# Heritability of leaf stable carbon isotope signature in a diversity panel of the C4 plant *Sorghum bicolor*

**DOI:** 10.64898/2026.08.16.745146

**Authors:** Joseph D. Crawford, Collin Luebbert, Ivan Baxter, Daniel P. Schachtman, Asaph B. Cousins

## Abstract

A strategy to improve agricultural water productivity is to increase water use efficiency (WUE) at the level of plant transpiration through genetic selection. This requires detectable genetic variability in WUE and the ability to phenotype and select plants with higher WUE within a population. A proxy for phenotyping leaf level WUE by measuring carbon isotope signature (δ^13^C_leaf_) has been supported by theory and data in C_4_ species. However, the functional relationship of δ^13^C_leaf_ and WUE in C_4_ species can be driven by genetics and environment. Therefore, a wide survey of existing natural variation is needed to quantify the heritability and identify various genetic factors that influence δ^13^C_leaf_ and WUE. In this study a genome-wide association panel was used to quantify the heritability of δ^13^C_leaf_. We measured δ^13^C_leaf_ across a population of 360 genetically diverse lines of the C_4_ species *Sorghum bicolor* with single nucleotide polymorphic (SNP) markers determined from whole-genome resequencing. This analysis was conducted on two independent field environments where heritability of δ^13^C_leaf_ was evident and was driven by small genetic effects from loci that were consistently identified across environments. Candidate genes are presented that offer insights on future targets to manipulate and explore the functional relationship between δ^13^C_leaf_ and *WUE*_i_ in C_4_ plants.

**Highlight:** A high-throughput tool to assess water use efficiency contains a strong heritable component in a C_4_ species

## Introduction

Agricultural water use is estimated to be only 45% efficient and one strategy to improve water productivity is to increase water efficiency at the level of plant transpiration (Hamdy *et al*., 2003). There is evidence that some crops may have been selected over time to be most productive through high water use when soil water is ample (Roche, 2015; Koester *et al*., 2016). As a result, these crops may not use water efficiently particularly when soil water availability is low. While this breeding strategy may have been viable under non-water limiting growth conditions, climate change is predicted to increase temperatures and alter precipitation patterns. Higher temperatures increase crop transpiration, the effects of which will be compounded in agro ecosystems receiving less water resulting in reduced agricultural production (H.-O. Pörtner, D.C. Roberts, E.S. Poloczanska, K. Mintenbeck, M. Tignor, A. Alegría, M. Craig, S. Langsdorf, S. Löschke, V. Möller, A. Okem (eds.)). Therefore, development of crop genotypes that are more water use efficient and resilient under drier conditions due to climate changes are needed (Condon *et al*., 2004). Improvements in water use efficiency at the plant level (*WUE*_plant_) hinges on the ability to phenotype genotypes with genetically improved water use efficiency within traditional breeding programs. Additionally, water use efficiency is likely not a fixed trait but variable under different environmental conditions. Therefore, phenotyping for water use efficiency must select for genetic variance in *WUE*_plant_ that is independent of environmental influences (Leakey *et al*., 2019). One component of *WUE*_plant_ that meets these requirements of being quantifiable and buffered from short-term environmental conditions is leaf-level productivity of photosynthesis per unit water lost (Ellsworth and Cousins, 2016).

A large proportion of *WUE*_plant_ is driven by rates of leaf transpiration, which is in part determined by genetically controlled rates of stomatal conductance (*g*_s_). At the leaf level, net rate of assimilation of CO_2_ (*A*_net_) relative *g*_s_ defines intrinsic water use efficiency (*WUE*_i_). In this measure, *g*_s_ is independent of vapor pressure deficit and so *WUE*_i_ becomes comparable across genotypes and variable environments (Ellsworth and Cousins, 2016). While the mechanics of *WUE*_i_ are well understood (von Caemmerer and Baker, 2007), the genetic factors required to breed or engineer higher *WUE*_i_ are only starting to be investigated in large genetically diverse populations (Gresset *et al*., 2014; Feldman *et al*., 2018; Ellsworth *et al*., 2020; Crawford *et al*., 2024). This is primarily hampered because phenotyping *WUE*_i_ with gas exchange instruments is slow and labor-intensive, restricting their use in plant breeding systems where high-throughput measurements are necessary.

An alternative to using gas exchange is to measure the ratio of ^13^C to ^12^C in dried leaf material (δ^13^C_leaf_). This method has been reported as a proxy for *WUE*_i_ in both C_3_ and C_4_ species (Farquhar *et al*., 1982; Farquhar, 1983; Henderson *et al*., 1998; Condon *et al*., 2004). Theory predicts that the carbon isotope composition integrated into leaf tissue would approximate *WUE*_i_ because of C_4_ biochemistry and diffusion (Farquhar, 1983). However, there is uncertainty in how well δ^13^C_leaf_ can predict *WUE*_i_ within a C_4_ species because i) there are potentially complex fractionations due to the CO_2_ concentrating mechanism (CCM); ii) C_4_ species have naturally low ranges of *g*_s_ and the ratio of intercellular and atmospheric CO_2_ which are key drivers of δ^13^C; iii) variation in the leakage rate of CO_2_ from the CCM; and iv) fractionations that discriminate against ^13^C but decouple it from *WUE*_i_ (Twohey *et al*., 2026). All these factors are integrated into δ^13^C_leaf_. But it is not understood if these factors are heritable within a C_4_ species.

Surveys of δ^13^C_leaf_ variation within C_4_ species have been done in small or genetically structured populations (e.g. bi-parental populations, advanced recurrent selection populations). This narrow diversity limits our understanding about the genetic contribution to δ^13^C_leaf_ variation at the species level (O’Leary, 1988; Hubick *et al*., 1990; Henderson *et al*., 1998; Cabrera-Bosquet *et al*., 2009; Gresset *et al*., 2014; Twohey *et al*., 2019; Ellsworth *et al*., 2020; Sorgini *et al*., 2021). The heritability of δ^13^C_leaf_ within a C_4_ species is also unclear because the genetic contribution relative to potentially strong environmental or sub-population influences are unknown. Therefore, a wider intraspecies analysis of the genetic control of δ^13^C_leaf_ is necessary to advance its use as a proxy for *WUE*_i_ in C_4_ species.

A suitable population to investigate the genetic control of δ^13^C_leaf_ in C_4_ species is the sorghum bioenergy association panel (BAP). The BAP was developed to dissect the genetic drivers of bioenergy traits through marker trait associations by including diverse accessions from global collections of sorghum representing the five sorghum races (Brenton *et al*., 2016; Songsomboon *et al*., 2021). The BAP contains genotypes with variable end uses (e.g grain, sweet, and high biomass) and demonstrates large phenotypic diversity. High biomass as a trait is distinct from grain yield and many past diversity panels aim to understand factors controlling grain yield. Additionally, individuals of the BAP were genotyped with SNP markers and the sorghum genome is sequenced and well annotated (Brenton *et al*., 2016; McCormick *et al*., 2018; Songsomboon *et al*., 2021). The phenotypic and genetic diversity of the BAP provides a platform to study the heritability of δ^13^C_leaf_ and overcome obstacles previous research populations have encountered.

In this study δ^13^C_leaf_ was surveyed in the genetically and phenotypically diverse population of C_4_ species *Sorghum bicolor* BAP in two field environments. This allowed us to test the hypothesis that δ^13^C_leaf_ is heritable within a C_4_ species and under the control of specific genetic loci. Estimates of δ^13^C_leaf_ heritability in sorghum, and genome-wide associations (GWAS) of δ^13^C_leaf_ to candidate genes with annotated function provide insights into factors driving the relationship of δ^13^C_leaf_ and *WUE*_i_ in sorghum.

## Materials and Methods

### Plant Materials and Field Sites

The BAP is a collection of diverse sorghum genotypes from the USDA Germplasm Repository Information Network and was constructed to facilitate discovery of genetic drivers of bioenergy traits (Brenton *et al*., 2016). The population’s conception and motivation was originally described in Brenton et *al*. (2016). The data presented here came from sampling the BAP population planted in two separate field experiments. The first location was in Clemson, SC (34.6834° N; 82.8374° W) in 2017 and the second in Scottsbluff, NE (41.8666° N, 103.6672° W) in 2019.

In South Carolina, the sandy loam soil was prepared with a custom pre-plant fertilizer 15(N)-18(P)-24(K) at 136 kg ha^-1^. The field was planted May 31, 2017 with 361 members of the BAP. The genotypes were planted in 3 x 3 m plots (4-rows per plot) at approximately 96,000 plants ha^-1^. A seed treatment that included a solution of Concep II, NipIt, Apron XL, and Maxim XL. Bicep II Magnum at a rate of 3.5 l ha^-1^ was applied as a pre-emergence herbicide. Atrazine was applied at 4.5 l ha^-1^ after planting but before any germination. The field was fertilized at a rate of 84 kg ha^-1^ layby N 30 days after planting but did not receive any supplemental irrigation. There were two applications of Sivanto SL insecticide at 515 ml ha^-1^ for sugarcane aphid control in July and August. On July 6-7, 2017, approximately 40 days after planting, the upper most fully expanded were sampled from two individual plants within each plot. The individual leaf tissue was measured for δ^13^C_leaf_ and values were pooled for each plot. The experiment was designed as 2 randomized complete blocks and so each genotype was represented as N = 2 biological replicates with two 2 subsamples per plot.

In Scottsbluff, Nebraska each plot was 1.52 m x 4.57 m planted with 2 rows spaced 76 cm apart and planted with 100 seeds per plot. The field design had 4 randomized complete blocks with 30 plots x 12 ranges grid. The blocks were 75 m long and 52 m wide. Two blocks were irrigated and two were scheduled to a declining soil moisture regime. However, leaves were sampled 4 days into a season-long differential irrigation treatment and so considered all 4 replicates to be the same and had no treatment effect. Two leaves from separate plants per plot were sampled and pooled for analysis such that n = 4 biological replicates per genotype.

### Weather Data

Weather data was retrieved from the National Oceanic and Atmospheric Administration Climate Data Online repository for SC and NE stations during the growing seasons: Clemson, SC station USC00381770, (2017) and Scottsbluff, NE station USC00257667 (2019).

### Carbon isotope signature of dry leaf material

Sampled leaves from the NE and SC field locations were cut and flattened in coin envelopes and dried at 65°C for 7 days. After drying, the leaves were punched with a custom punch directly into tin capsules (Ellsworth *et al*., 2017) (Costech, Valencia, CA, USA). The punches were combusted into N_2_ and CO_2_ in an elemental analyzer (ECS 4010, Costech Analytical, Valencia, CA). The combusted gases were separated in a 3 m gas chromatography column and the isotope signature of the gases are measured with a continuous flow isotope ratio mass spectrometer (Delta PlusXP, Thermofinnigan, Bremen). Isotope ratios are reported per mill relative to Vienna Peedee belemnite standard (VPDB). Samples were measured in the Washington State University Stable Isotope Core Laboratory.

### Calculation of heritability

Heritability was calculated with a mixed model in the lme4 R package and then extracting the variance components from the model:

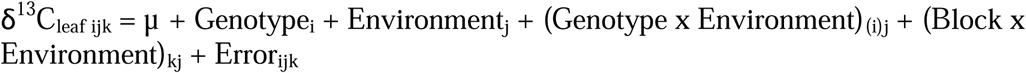

where δ^13^C_leaf ijk_ is the result of the mean (µ), i^th^ Genotype, j^th^ Environment, the interaction of the i^th^ genotype nested within the j^th^ environment, the k^th^ block nested in the j^th^ environment, and the residual error. The number of replicates differed between the two locations so the harmonic mean number of non-zero replicates of genotype entries was used as an adjustment in the calculation of broad sense heritability for multiple environment trials (Holland *et al*., 2002):

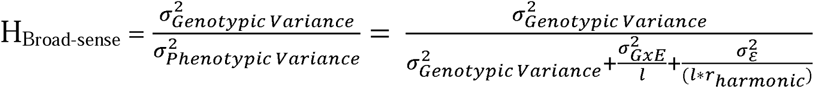

where 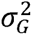 is genotype variance component, 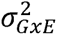 is the genotype by environment variance component, 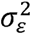 is the the residual variance, l is the number of independent environments, and T_harmonic_ is the harmonic mean number of replicates calculated as the average, non-zero occurrences of a genotypes across environments. Bootstrap resampling of the heritability estimate was accomplished by sampling with replacement entire blocks of the data set for 50,000 simulations. By sampling entire blocks the design structure and covariance of the real experiment are maintained. H^2^ was calculated for each simulation and the mean, and 95% confidence intervals were calculated across all simulations.

### Genetic markers of the Bioenergy Association Panel (BAP)

A total of 390 genotypes within the bioenergy association panel were resequenced at the HudsonAlpha Institute for Biotechnology with 150-bp paired-end short end Illumina X10 reads as part of the TERRA-REF project (Songsomboon *et al*., 2021). Single nucleotide polymorphic (SNP) markers were subset from whole-genome resequencing of 390 originally individuals included in the BAP. The raw reads are available at DataCommons.cyverse.org. This data set was larger than needed for GWAS so it was further reduced in size by taking SNPs from chromosomes 1-10 and 1 SNP per approximately every 2,000 bp using the htslib 1.8, bcftools 1.10.2, and TASSEL 5.2.67 libraries and the following BASH script:

- bcftools view BAP_new.recode.vcf.gz -r Chr01,Chr02,Chr03,Chr04,Chr05,Chr06,Chr07,Chr08,Chr09,Chr10 -Oz -o BAP_1_10_Full.vcf.gz
- bcftools +prune BAP_1_10_Full.vcf.gz -w 2000bp -n 1 -Ov -o BAP_1_10_reduced_2000bp_marker.vcf

The output was converted from vcf into HapMap format using TASSEL:

- run_pipeline.pl -Xms64G -Xmx64G -fork1 -vcf BAP_1_10_reduced_2000bp_marker.vcf
- - export BAP_1_10_2kbp_marker -exportType Hapmap

### Clustering Subpopulations

The BAP racial class identifiers were incomplete within the dataset. Therefore, for each genotype the genetic distance matrix between individuals in the BAP derived from SNP markers was used to cluster the panel into 10 unequally sized groups via unweighted pair grouping with arithmetic mean (UPGMA) hierarchical clustering.

### Genome-Wide Association with Compressed Mixed Linear Model

GWAS of δ^13^C_leaf_ in the BAP was performed with GAPIT2 and GAPIT3 software (Lipka *et al*., 2012; Tang *et al*., 2016; Wang and Zhang, 2021). GAPIT functions were sourced from https://zzlab.net. The compressed mixed linear model (CMLM) was used with one and/or two principal components to control for population structure (Zhang *et al*., 2010). The K matrix (kinship or relatedness matrix) was calculated within GAPIT with the default VanRaden method (VanRaden, 2008). The compression of groups was achieved with the method that takes the mean of clusters and mean group kinship. Markers were filtered to have a minor allele frequency more than 5%.

### Multivariate Adaptive Shrinkage (MASH) of SNP Marker Effects

GWAS was performed using CMLM on each of the complete blocks within the outdoor experiment environments (Nebraska and South Carolina locations) and considered these separate for the analysis (total N =6). After extracting marker effects with CMLM from each of these six sites, the multivariate adaptive shrinkage (MASH) method was used to identify strong effect SNPs across environments with the reasoning that strong SNPs would be significant in many or all environments, although many could be insignificant by chance in others. Urbut et al.’s (2019) implementation of MASH in R from CMLM GAPIT output was achieved through wrapper functions gapit2mashr which are available on Github (https://github.com/Alice-MacQueen/gapit2mashr) (MacQueen *et al*., 2020; Lovell *et al*., 2021). The final effect sizes are posterior probabilities. Marker significance is on a log10 Bayes Factor scale.

### Candidate Gene Analysis

An R package, PanvaR, was used to refine the likelihood of candidate causal genes within an interval of GWAS SNP hits (Luebbert *et al*., 2026, Preprint). Briefly, PanvaR calculates local linkage dis-equilibrium for the genomic region for which a significant QTL from GWAS and MASH was found. Then the software re-runs GWAS with a single-locus generalized linear model (GLM) with gene models in context and evaluates a SNP’s impact with a PlantCaduceus score. The PlantCaduceus zero-shot score comes from a DNA language model that predicts causal variants across 16 angiosperm genomes (Zhai *et al*., 2025). PanvaR also includes alternative scores of prediction that a gene model is causal such as the p-value of the local GLM GWAS, the estimate of linkage disequilibrium between the marker and the gene model, and the weighted average of all the scores (snp.score).

### Statistical Analysis

Statistical analysis of sorghum genetic clusters for differences in δ^13^C_leaf_ was assessed with an analysis of variance test. The clusters were formed with unweighted pair grouping with arithmetic mean (UPGMA) hierarchical clustering. Statistical analysis was done with R version 4.3.1 (R Core Team, 2018).

## Results

### Growing Season Weather Data

In Clemson, SC the mean daily maximum temperature from planting until the day of sampling was 28.8 °C with a range from 16.1 to 33.9 °C. The mean daily precipitation was 5.19 mm with a range from 0 to 47.8 mm. In Scottsbluff, NE the mean daily maximum temperature from planting until the day of sampling was 28.0 °C with a range from 13.9 to 36.7 °C. The mean daily precipitation was 2 mm with a range of 0 to 15 mm.

### Heritability, and genetic variation of ***δ***^13^C_leaf_

In Clemson, SC the BAP panel was grown in two field randomized complete blocks. Across all genotypes, the mean δ^13^C_leaf_ value was -13.23‰ with a standard deviation of 0.18 ‰ (Fig 1). The spread in δ^13^C_leaf_ was 1.15 ‰, with maximum and minimum values between -12.67 and -13.83 ‰ (Fig 1). The range of δ^13^C_leaf_ in this study is 2-fold higher than two reports with 12 or 30 sorghum accessions of field-grown plants (Hubick *et al*., 1990; Henderson *et al*., 1998), and 2.8-fold higher than a greenhouse experiment with 30 genotypes, (Henderson *et al*., 1998). There were 361 genotypes whose seeds germinated, plants established, and were sampled for δ^13^C_leaf_.

**Fig 1.**
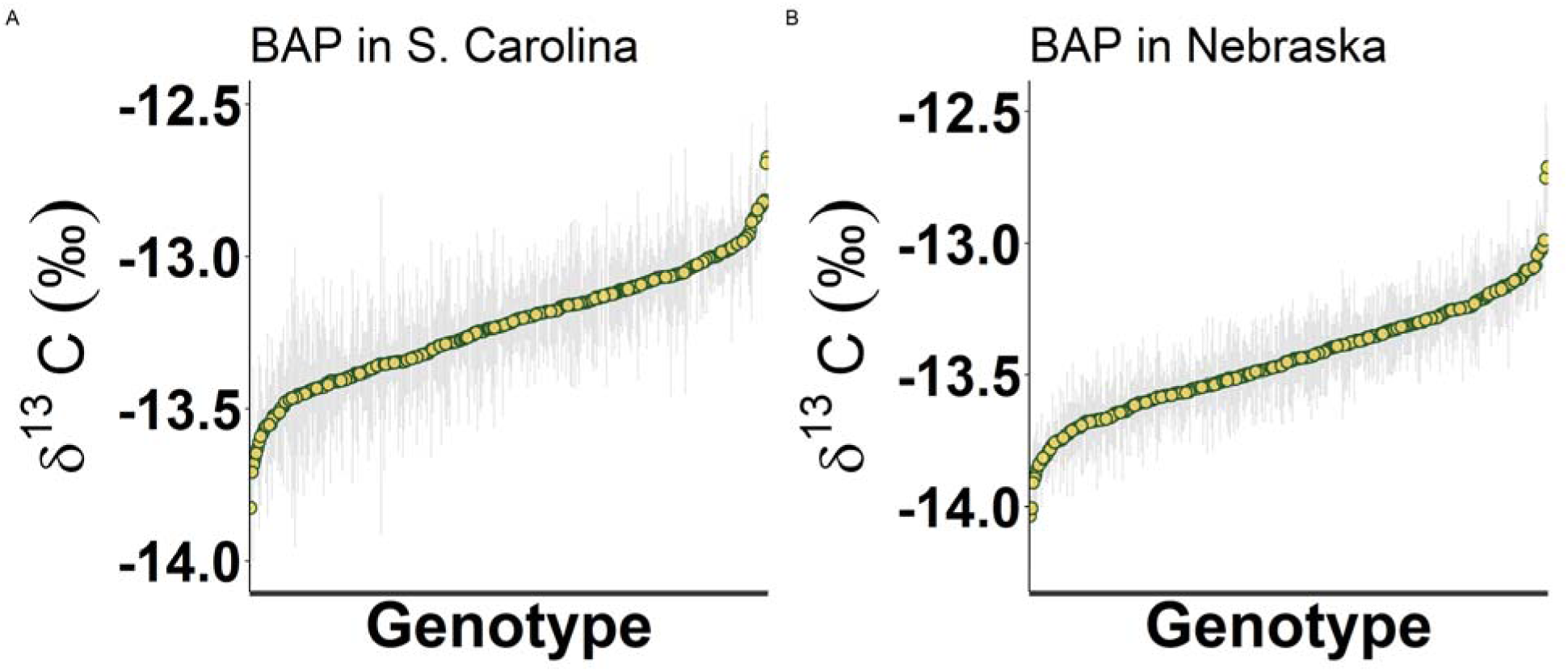
δ^13^C_leaf_ signatures of sorghum genotypes from the Bioenergy Association Panel (BAP) grown in two environments ranked from lowest to highest. A) Clemson, South Carolina location. B) Scottsbluff, Nebraska. Points represent one genotype and error bars represent standard error of the genotype mean. N =2 in SC and N = 4 in NE. The δ^13^C_leaf_ leaf from these locations comes from separate years but in both years δ^13^C_leaf_ was sampled approximately 40 days after planting.

In Scottsbluff, NE the BAP was under two field treatments that consisted of two irrigated complete blocks (WW), and two complete blocks where the irrigation schedule was gradually reduced to achieve a water stressed condition (WS) relative to WW. However, samples were collected early in the season before the differential water regime had an any impact on soil moisture and there was not a significant treatment effect when using the two blocks to test for a treatment effect on δ^13^C_leaf_ (ANOVA; p-value = 0.839). Hereafter, the four blocks were regarded as replicated complete blocks without a treatment. In total there were 363 genotypes whose seeds germinated, plants established, and were sampled for δ^13^C_leaf_. Across all replicates and genotypes the mean was -13.4 ‰ and the standard deviation was 0.2 ‰ (Fig 1). The spread was 1.3 ‰ with maximum and minimum values between -12.7 and -14.0 ‰ (Fig 1).

The broad sense heritability of δ^13^C_leaf_ for the BAP population, defined as the proportion of phenotypic variance explained by genotype, was 57.5%. Bootstrap resampling of phenotypes by resampling block level data demonstrated a mean bootstrap broad-sense heritability of 48%, and a 95% confidence interval from 37% to 57.5%. The environmental variance component accounted for 16% of the total model variance. GxE variance was 3.3% of the total model variance.

### Subpopulations Clustered with Genetic Markers Have Significant Differences in ***δ***^13^C_leaf_

The influence of sorghum races or subpopulations on δ^13^C_leaf_ was statistically evaluated. The BAP includes the five major races of sorghum, intermediates, and admixtures (Brenton *et al*., 2016). However, instead of using these racial class identifiers for each genotype, the genetic distance matrix between individuals in the BAP derived from SNP markers was used to cluster the panel into 10 unequally sized groups via unweighted pair grouping with arithmetic mean (UPGMA) hierarchical clustering (Fig 2). There were significant differences in δ^13^C_leaf_ between groups in the NE and SC sites (Fig 2; ANOVA; 9 d.f.; NE p-value = 5.4E-14; SC p-value = 5.7E-6). The groups were mostly consistent in their rank order trends of δ^13^C_leaf_ values between NE and SC experiment sites except for cluster 10. For example, δ^13^C_leaf_ in groups 3 and 5 were greater than the mean. Groups 2 and 9 were consistently more negative (depleted) than the mean (Fig 2). Small groups, such as group 10, changed rank from first to last between sites but contained only 1 genotype (Fig 2).

**Fig 2.**
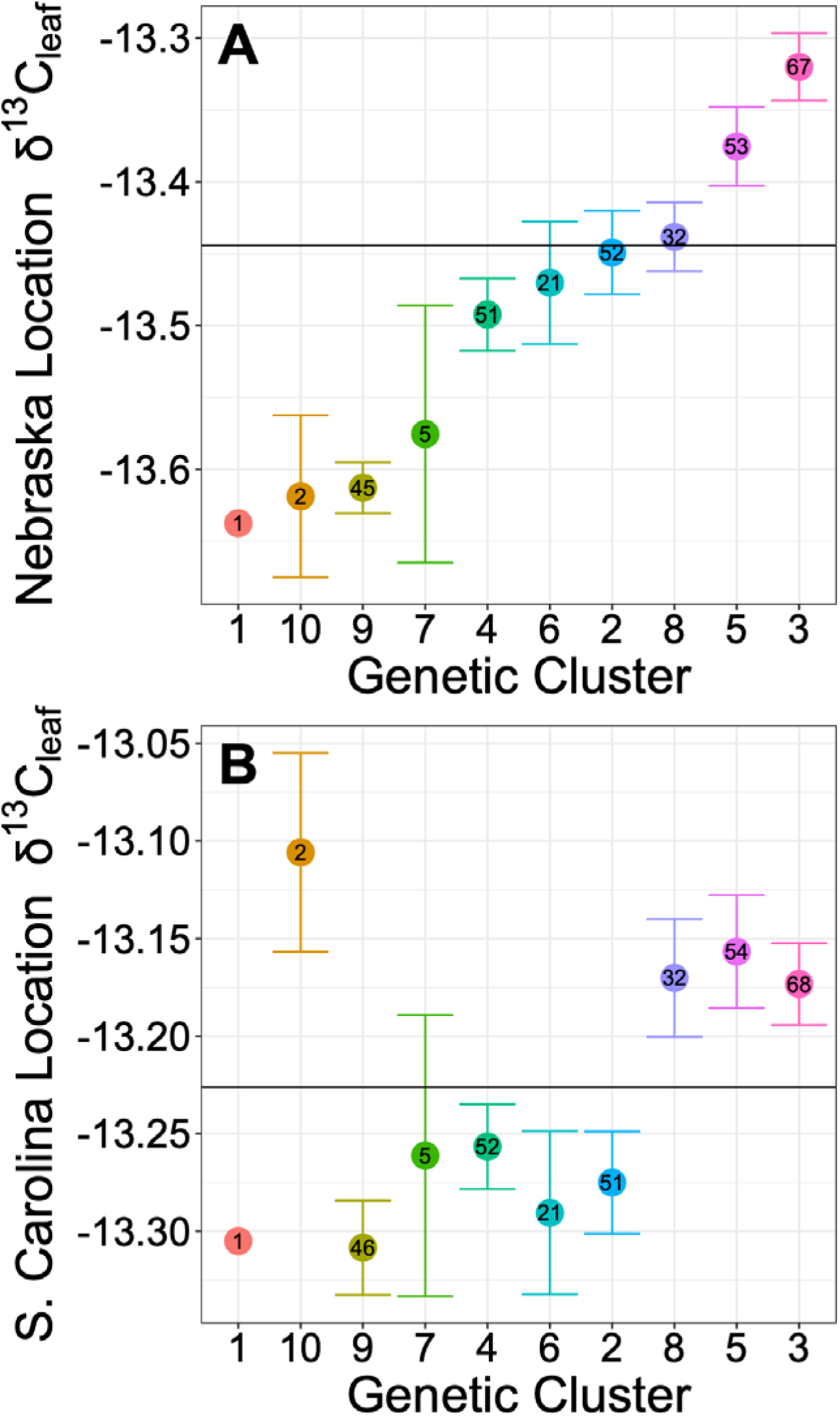
Means and SE of the δ^13^C_leaf_ of genetic groups of the Sorghum Bioenergy Association Panel (BAP) that were classified by calculating genetic distance with SNP markers and clustering into 10 groups using hierarchical clustering. A) Results from NE in outdoor field conditions. B) Results from SC in outdoor conditions. The clusters are ordered on the x-axis by δ^13^C_leaf_. The black line is the grand mean of all genotypes for each location. Genetic groups show similar rank-order trends between locations. Rank switching of group 10 is likely statistical noise and only contained two individuals.

### ***δ***^13^C_leaf_ and Genetic Marker Derived Principal Components (PC)

Subpopulations within the BAP were examined with principal components derived from SNP markers. The first PC explained 16% of the variance. The second and third PC explained 9 and 6%, respectively. PC1 grouping of genotypes was similar to groups organized with hierarchical clustering as evidenced by the similarity of cluster member’s order along the range of PC1 (Fig 3). The relationship of PC’s (as a proxy for population structure) and δ^13^C_leaf_ with linear regression demonstrated there was no significant relationship between PC1 and δ^13^C_leaf_ (Fig 3; regression; p-value = 0.42).

**Fig 3.**
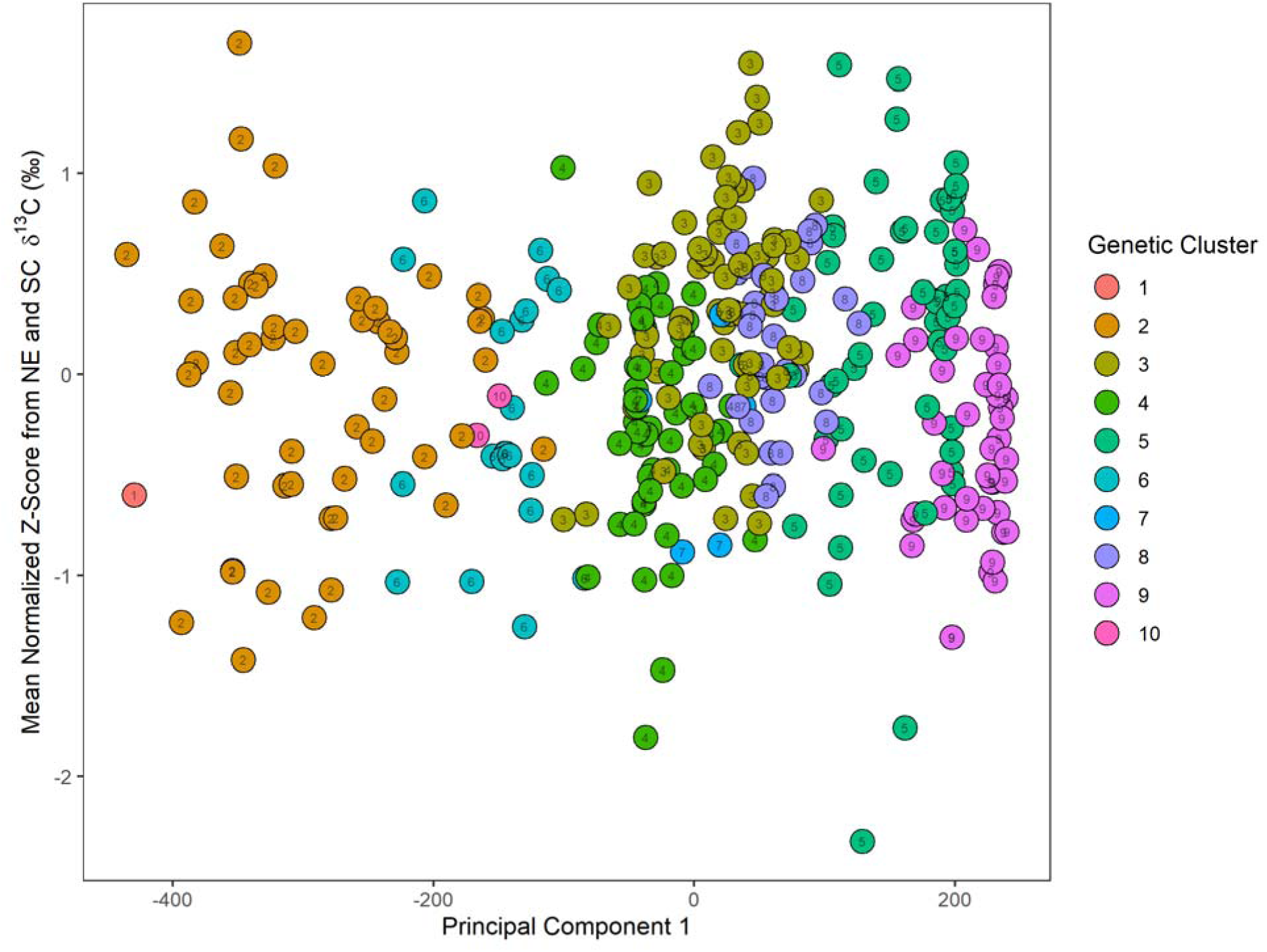
The first principal component of the genetic marker data set, representing a genomic region that explains the largest percentage variation between genotypes in the BAP, plotted against the normalized δ^13^C_leaf_ of genetic groups. There was no significant linear relationship between δ^13^C_leaf_ and PC1. The clusters in colored points, determined via hierarchical clustering in parallel, fall along the PC1 axis and appear to coalesce into similar clusters.

Additionally, genotypes were annotated with previously-known classifiers such as their photoperiod sensitive/insensitive phenotype and their production use-type (sweet, grain, or cellulosic sorghum) within sub-populations from hierarchical clustering (Brenton *et al*., 2016). Unfortunately, each use-type was not always represented in high frequency within groups. There was no significant effect from the group’s photoperiod phenotype (ANOVA; p-value = 0.84; data not shown) or use-type (ANOVA-p-value = 0.73; Fig 4).

**Fig 4.**
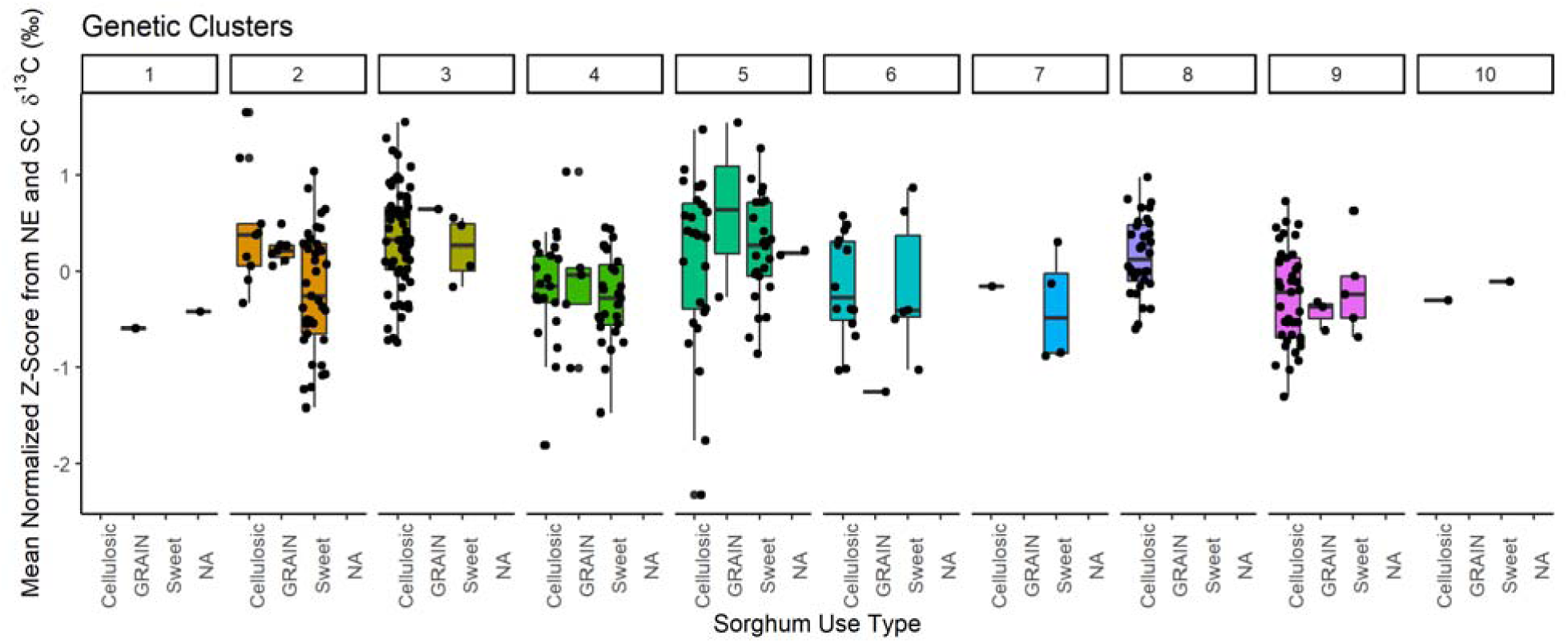
Normalized Z-score δ^13^C_leaf_ boxplots of BAP genetic clusters in facets, categorized for Use type: grain, sweet, cellulosic. There was no significant effect from use-type. Use type was not represented in each genetic cluster and for some genotypes there was no labeled use type (NA). Cellulosic sorghum genotypes have higher representation in the BAP. Line within boxplots represents the median and box upper and lower boundaries represent the 25 and 75% quartile. Points are plotted atop boxplots to show representation numbers.

### Genome-wide association finds repeatable SNPs across replicates that impacts ***δ***^13^C_leaf_

The sorghum genome was scanned for significant associations between 300,165 SNP markers and a genotype’s δ^13^C_leaf_ signature in the SC and NE environments. The sorghum genome is estimated to be 351 Mb (Phytozome.org), therefore it was estimated the genome was saturated with a genetic marker approximately every 1,170 bp. The resolution of detection was limited by the linkage disequilibrium which was estimated to be 15-150 kbp in the BAP (Brenton *et al*., 2020). There were 337 genotypes out of 360 that had no missing data components (e.g. genotype markers, phenotype data, and PCs necessary for GWAS calculations across the two environments. To identify SNPs that robustly impact δ^13^C_leaf_, consistent SNP signals that occurred across environments were identified. Each randomized complete block within each outdoor environment (SC and NE) was considered as an independent environment and performed GWAS on the δ^13^C_leaf_ phenotype with CMLM. To further examine the genome-wide distribution of SNP’s with a signal in more than one environment the entire SNP marker p-value and effect sizes data from each environment was combined and evaluated with multivariate adaptive shrinkage (MASH) (Urbut *et al*., 2019; Lovell *et al*., 2021). Across the 6 environments there were 2 SNPs that were significant in all 6 environments and present on chromosomes 1 and 9 (Table 1; Fig 5). The SNP Chr09_50221901 effect size ranged from 0.0173 to 0.0675 ‰ across the replicates and the mean was 0.0411 ‰. The SNP Chr01_21911032 effect size ranged from 0.0235 to 0.0837 ‰ across the replicates and the mean was 0.0495 ‰. The strongest PlantCaduceus scores within 100 kilobases of the snps in chromosomes 9 and 1 were 9.23 and 9.33, respectively (Table 1; Supplementary Tables 1;2; Note: the signs of the zero-shot score are reversed in PanvaR for convenience). Using 2 PCs reduced the discovery of significant SNPs to the one on chromosome 9.

**Fig 5.**
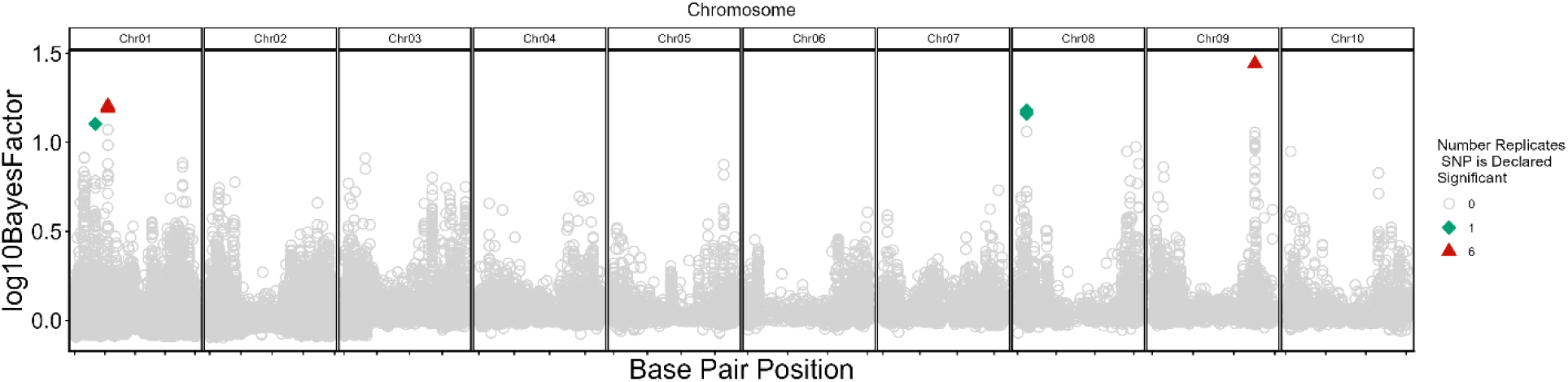
GWAS of δ^13^C_leaf_ across 6 environments finds significant SNPs with strong effects in more than one environment. GWAS was performed on the phenotypes from each of the 6 environments with the compressed mixed linear model (CMLM). In this analysis the 6 environments were the randomized complete blocks in two outdoor environments (4 blocks NE; 2 blocks SC). The marker effects were extracted from those analyses and compiled as input into multivariate adaptive shrinkage (MASH). MASH finds strong effect SNP markers across environments relative to SNPs expected to have null effects. The x-axis within chromosome facets represents physical base pair position. The strength of association is determined on a log10 Bayes factor scale. Point shape provides information on the number of environments (out of 6 environments) the SNP was declared significant in.

**Table 1.** The most significant SNP GWAS markers for δ^13^C_leaf_ and gene models with potentially protein altering variants.

| SNP Marker | Chromosome | Physical Position | MASH Effect Size Across Environments (‰) | Corresponding Gene Models in LD<sup>†</sup> with the SNP and found significant from PanvaR<sup>‡</sup> | Functional annotations from Arabidopsis |
| --- | --- | --- | --- | --- | --- |
| Chr01_21911032 | 1 | 21911032 | 0.03±0.014 | Sobic.001G227800 | DUF1338 domain-containing protein |
|  |  |  |  | Sobic.001G229000 | xyloglucan galactosyltransferase MUR3, putative, expressed |
| Chr09_50221901 | 9 | 50221901 | 0.03±0.014 | Sobic.009G145000 | Early-Responsive To Dehydration Stress/ OSCA1 |
|  |  |  |  | Sobic.009G144600 | Peroxidase / Lactoperoxidase |

<sup>†</sup>Linkage Disequilibrium <sup>‡</sup>PanvaR is a software program that predicts which gene models are influencing the phenotype in regions around a SNP marker

### Combinations of additive alleles promoting/reducing ***δ***^13^C_leaf_ occur in low frequency within the BAP

Genotypes were examined for their different allelic combinations of the 2 strongest SNP markers discovered from GWAS and MASH and their frequency in the BAP. For a marker, each allele was classified as the high (H) or low (L) depending on the allele’s mean δ^13^C_leaf_ transformed Z-scores was greater or less than zero. There were 4 combinations found in the BAP after excluding individuals that were heterozygous at any marker. The highest frequency GWAS QTL combination was present in 35% of BAP genotypes and carries LL, a negative-Z-scored allele for each of the 2 markers displayed). Generally, the δ^13^C_leaf_ phenotype on a Z-score scale for each genotypic allele combination decreased with more L alleles present and increased with more H alleles (Fig. 6).

**Fig 6.**
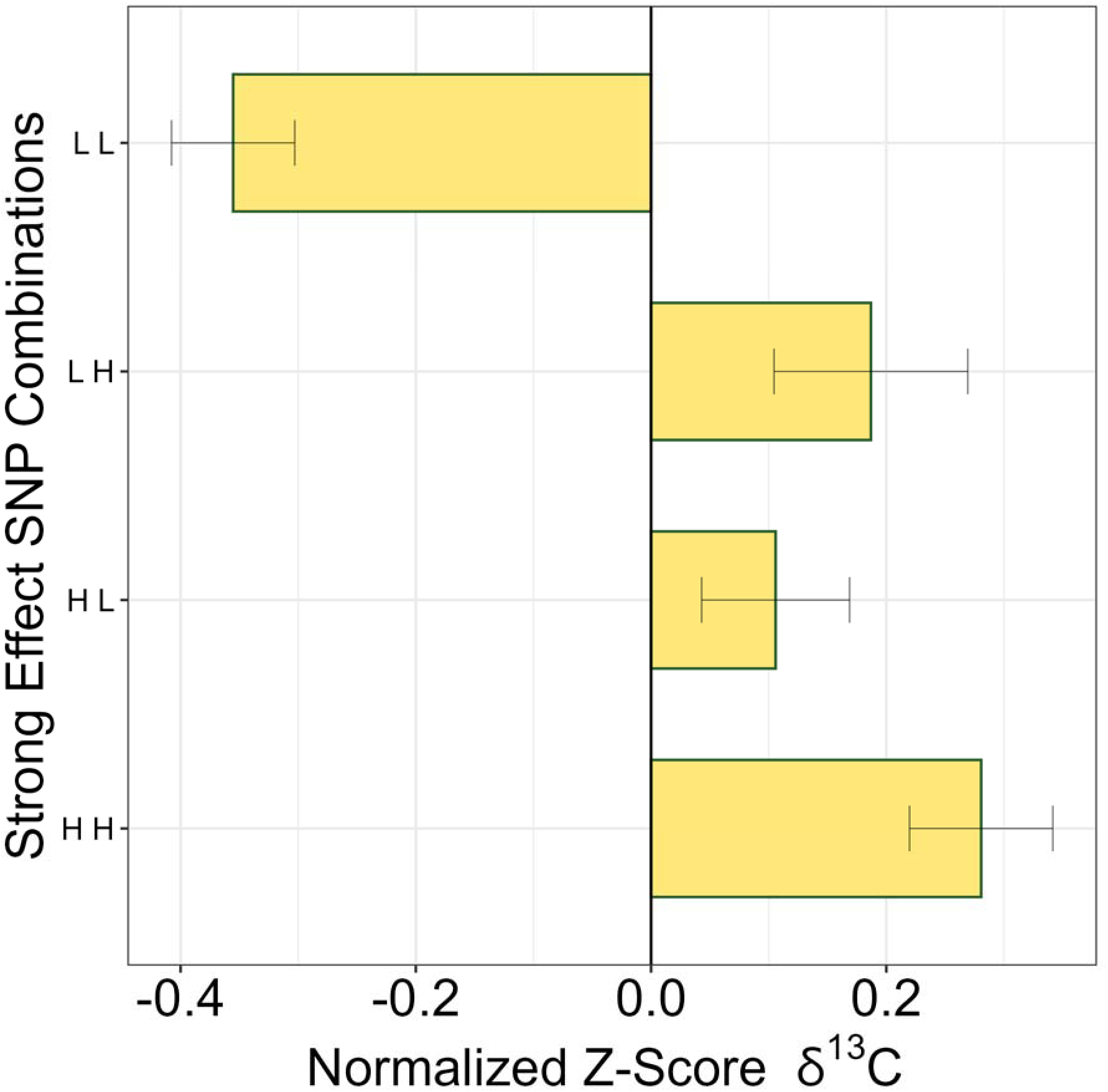
Allelic combinations carried by genotypes of the 2 strongest SNPs associated with δ^13^C_leaf_ identified from GWAS and MASH. See Table 2 for specific allele information. The existing allele combinations in the population were identified and their Z-score normalized δ^13^C_leaf_ signature from outdoor environments was summarized as a Z-score mean and standard error.

**Table 2.** Allelic combinations of strong effect GWAS QTL and phenotypic effects.

| First Strong QTL SNP<br>*Chr09:50221901* | Second Strong QTL SNP<br>*Chr01:21911032* | Allele Combinations | Phenotypic Effect | Sample Size (n) | Proportion of diversity panel |
| --- | --- | --- | --- | --- | --- |
| A = Low | C = Low | AC | Low–Low (LL) | 118 | 35% |
| A = Low | T = High | AT | High–Low (HL) | 60 | 18% |
| G = High | C = Low | GC | High–Low (HL) | 60 | 18% |
| G = High | T = High | GT | High–High (HH) | 90 | 27% |

## Discussion

The genetic control of leaf level water use efficiency (*WUE*_i_) in C_4_ plants is largely unknown because of a lack of a high-throughput measurement to survey broad phenotypic diversity. The ratio of ^13^C to ^12^C in dried leaf material (δ^13^C_leaf_) has the potential to approximate *WUE*_i_ in C_4_ plants but there is uncertainty on how much δ^13^C_leaf_ is influenced by genetics compared to environmental conditions. The goal of this study was to estimate the heritability of δ^13^C_leaf_ within a diverse C_4_ species population and to discover potential candidate genetic loci influencing δ^13^C_leaf_. The results demonstrate that δ^13^C_leaf_ is heritable in a diverse sorghum population and genetic associations with δ^13^C_leaf_ provide insights into factors driving the relationship of δ^13^C_leaf_ and *WUE*_i_ in this C_4_ species.

### Variation of ***δ***^13^C_leaf_ across the *S. bicolor* bioenergy association panel (BAP)

The range of δ^13^C_leaf_ in sorghum in this study (1.3 ‰) was similar between the two outdoor common field experiments (Fig 1). The range was about double the previous reports for sorghum. For example, a survey of 12 sorghum genotypes found a range of 0.6 ‰ in field grown plants (Hubick *et al*., 1990), and another study found a range of 0.41 and 0.67 ‰ in a greenhouse and field experiment with 30 genotypes, respectively (Henderson *et al*., 1998). One potentially important difference between the current study and previous reports is that grain sorghum genotypes are a small proportion of total plants in the BAP (Fig 4), whereas in previous reports the genotypes were almost exclusively grain sorghums (Hubick *et al*., 1990; Henderson *et al*., 1998). A hallmark of artificial selection in grain relative to landrace sorghum is pre-programmed terminal growth upon flowering (photoperiod insensitivity) (Hao *et al*., 2021). This has important implications because sorghum genotypes that flower later potentially have prolonged genetically and environmentally-dependent physiology that can influence δ^13^C_leaf_ differentially over time. This was suggested by Hubick *et* al. (1990) when they found a stronger correlation between δ^13^C_leaf_ and sorghum grain yield in their analysis of two late-flowering genotypes that endured late season water stress. However, there was no evidence that grain sorghum intrinsically influenced δ^13^C_leaf_ in our study (Fig 4). The greater range of δ^13^C_leaf_ in the BAP reported here is likely related to the relatively large genetic diversity of this population. However, the δ^13^C_leaf_ values of the BAP are within ranges previously seen in other C_4_ species. For example, mean differences between *Setaria italica* and *S. viridis* was 2 ‰ and varieties of *Zea mays* differed by about 1.5‰ (Twohey *et al*., 2019; Ellsworth *et al*., 2020). It is notable that the *Setaria* experiment used a biparental population from an intraspecific cross which strongly contrasted in many traits (Devos *et al*., 1998). Additionally, the *Z. mays* data cited here was from diverse maize landraces along with some of their outcross progeny (Kolbe *et al*., 2018; Twohey *et al*., 2019). Taken together, the genetic diversity within a large population like the BAP should be sufficient to test for δ^13^C_leaf_ heritability.

Broad-sense heritability, defined as the variance of the phenotype attributable to a genotype relative to the total phenotypic variance, of δ^13^C_leaf_ was 57.5% in the BAP population. In turn, 42% of the phenotype variance is influenced by non-genetic factors. There are a few reasons that can explain non-genetic factors of δ^13^C_leaf_ in this population. The first is that δ^13^C_leaf_ is sensitive to environmental factors like water stress. The theory for leaf carbon isotope discrimination (Δ^13^C) during C_4_ photosynthesis predicts stomata closure will shift δ^13^C_leaf_, and this effectively increases the environmental variance which can influence estimates of heritability. For example, in the biparental population of *Setaria,* the heritability of δ^13^C_leaf_ was 50% under a well-watered treatment but reduced to only 5% under a water-stress treatment (Ellsworth *et al*., 2020). Estimates of heritability can also be population specific and sensitive to which genotypes are included in an experimental population, similar to the range effect on δ^13^C_leaf_ discussed above (Dohm, 2002). Advanced experimental design to ameliorate and estimate block and environment effects should be prioritized in future work. The overall heritability of δ^13^C_leaf_ in C_4_ sorghum can become clearer by testing in more environments but it is likely that there are substantial environmental influences on δ^13^C_leaf_ which might interact with specific characteristics of genetic sub-populations.

### ***δ***^13^C_leaf_ signature in sub-populations

Sorghum sub-populations (Murthy and Govil, 1967; Brown *et al*., 2011; Songsomboon *et al*., 2021) had distinguishing δ^13^C_leaf_ signatures within the BAP (Fig 2). An alternative way of summarizing and reducing the genetic complexity of the population was the principal components of the genetic markers for the population. The genetic clusters classified via hierarchical clustering roughly fell along the PC1 axis indicating agreement between the two methods (Fig 3). Regardless of approach, the differences in δ^13^C_leaf_ between clusters is likely due to differences in allele frequencies of causal genes for δ^13^C_leaf_ between clusters. However, regression analysis did not find a statistically significant relationship between PC1 and δ^13^C_leaf_ (Fig 3). Taken together, this shows that δ^13^C_leaf_ differs between clusters, but δ^13^C_leaf_ does not *define* these genetic clusters. This is important because it indicates genetic cluster and δ^13^C_leaf_ phenotype are not confounded which could lead to misleading marker-trait associations. The hypothesis that δ^13^C_leaf_ was influenced by important genotype categorizations such as photoperiod sensitive/insensitive and use-type (grain, sweet, or cellulosic) was also tested, and there was no impact of photoperiod or use-type classification on δ^13^C_leaf_ (Fig 4). This is important because certain compounds like sucrose and starch have different isotopic signatures through post-photosynthetic fractionations and accumulation of specific carbon compounds in sweet sorghums could shift δ^13^C_leaf_ (Tcherkez *et al*., 2011; Caemmerer *et al*., 2014). In the absence of any major confounding correlations genome-wide association was used to identify potential genetic variants influencing δ^13^C_leaf_.

### Genetic variants drive variation in ***δ***^13^C_leaf_

Genome-wide association (GWAS) discovered small effect loci associated with δ^13^C_leaf_ (Fig 5; Fig 6; Table 1). The potentially large environmental influence on δ^13^C_leaf_ motivated an effort to demonstrate the repeatability of detecting causal SNPs for δ^13^C_leaf_ across several environments. Multivariate adaptive shrinkage (MASH) was used to find correlations of SNP markers effects between replicates and build a model to determine the null and strong effect markers, and to identify SNPs that had strong signals across environments (Fig 5). With MASH, the number of markers deemed significant was influenced by which principal components were added to the GWAS mixed model. Adding 2 PCs reduced significant SNPs to 1, but it was a subset of the larger set discovered with 1 PC. Statistical significance of a marker in all blocks is considered strong, reliable evidence of their association (Brenton *et al*., 2020; MacQueen *et al*., 2020; Lovell *et al*., 2021).

It is not surprising that the genetic architecture of δ^13^C_leaf_ in this C_4_ species appears to be driven by small effect loci since it is a quantitative trait subject to potentially strong environmental influence (Ellsworth *et al*., 2020). Therefore, differences in δ^13^C_leaf_ across this population may in part be explained by the quantitative differences in several SNP allele combinations within individual genotypes. Assuming additive allelic effects, these small effect loci can confer a large range of phenotypic variation at the population level (Fig 1), and when stacked within a given genotype (Fig 6). This stacking of alleles suggests that entire populations could achieve higher or lower δ^13^C_leaf_ through selection or marker-assisted selection in a breeding population. Indeed, combinations of the 2 significant markers (Table 1) had an additive effect on δ^13^C_leaf_, similar to the trend in *Setaria* where additive alleles correlated with δ^13^C_leaf_ (Ellsworth *et al*., 2020). Insights into the functional role of candidate genes under the significant markers on δ^13^C_leaf_ could provide evidence for their potential role in *WUE*_i_.

### Candidate genes

In this BAP population two SNPs were identified as the strongest effects that were repeated in all six replicates of the outdoor experiments and are expected to be the highest confidence loci that putatively impact δ^13^C_leaf_ (Table 1). The SNP Chr09:50221901 had two gene models within its region of LD that were predicted via PanvaR to be causally modifying δ^13^C_leaf_ phenotypes (Table 1). The gene model annotations under the peak suggest that it is a functional genomic region (Supplementary Table 1). The gene model with the strongest PlantCaduceus causal prediction score was a class III peroxidase gene model (Sobic.009G144600) repeated 7 times with the copies surrounding an Early-Responsive to Dehydration (ERD) family gene (Sobic.009G145000). Both are localized in guard cell protoplasts in *Arabidopsis* (Obulareddy *et al*., 2013).

Previously, the ERD gene family was characterized as integral membrane proteins with domain-of-unknown-function 221 (DUF221). Several genes within the family have demonstrated roles in abiotic stress, but most notably drought stress, and have been shown to control sensitivity to abscisic acid-promoted functions (Kiyosue *et al*., 1994; Kariola *et al*., 2006). More recent work has reclassified ERD genes in plants as reduced-hyperosmolality-induced-[Ca^2+^]_i_-increase-1 (OSCA1). Through osmotic stress screening, *OSCA1* was found to be mediating hyperosmolality-induced calcium increases at the plasma membrane and characterized as an osmotic sensor (Hou *et al*., 2014; Yuan *et al*., 2014). Fascinatingly, OSCA1 has more broadly been described as a mechanically gated ion channel that converts mechanical tension of membrane lipids into electrical signals (i.e. channel opening) (Jojoa-Cruz *et al*., 2018; Murthy *et al*., 2018; Zhang *et al*., 2018; Deng *et al*., 2026) . Therefore, more generally, OSCA1 has the potential to be a mechanosensitive channel that directly transduces hydraulic signals such as changes in water potential by sensing membrane tension or turgor (Scharwies and Dinneny, 2019). This particular gene model annotation OSCA1.7 in *Arabidopsis* (AT4G02900.1) has been shown to be independent of the abscisic acid stomatal closure pathway (Thor *et al*., 2020; Yoshioka and Moeder, 2020).

Nunes et al. (2023) identified a *Brachypodium distachyon* class III peroxidase mutant with reduced prickle hair size, longer stomata, increased stomatal conductance, decreased *WUE*i, and disrupted pavement cells near stomata with no changes in stomatal density. The resulting stomatal elongation also disrupted the well-established negative correlation of stomatal size vs density. They hypothesized that stomatal elongation balances the disruption of mechanical forces (e.g. turgor) of the epidermis when prickle hair cells are reduced by the peroxidase (Nunes *et al*., 2023). This could incorporate the mechanical sensing of pavement cell forces by guard cells via variance in OCSA1 thresholds in sorghum. The role of the 7 peroxidases could indicate that they are involved with hydrogen peroxide-mediated stomatal activity (Pei *et al*., 2000; Shi *et al*., 2024), cell wall stiffening by lignification (Hatfield *et al*., 2017; Hu *et al*.), or limiting cell wall expansion (Raggi *et al*., 2015). Crawford et al. (2025) reported that under steady-state conditions, stomatal aperture was not uniform in maize leaves, however this observation raises the question of how guard cells might achieve variance in aperture despite homogeneous turgor/signaling conditions. These candidate genes offer some insight into that mechanism, where global leaf turgor is uniform, aperture size is relative to mechanical strength of the guard cell walls which could be modified with altered proportions of polysaccharide compounds (e.g., xylan, cutin, pectin). Indeed, the gene model with the second highest PlantCaduceus prediction score was tandemly duplicated Trichome Bi-Refringence-like 7 gene models (Sobic.009G143900; Sobic.009G144050; Supplementary Table 1) and are involved with secondary cell wall maintenance probably through altering pectin or xylan, although these gene’s specific functions are not well understood (Bischoff *et al*., 2010).

The analysis of the second significant marker (chromosome 1:21911032) follows a parallel pattern (Supplementary Table 2). Using PanvaR’s snp.score metric, the most likely gene is a xyloglucan galactosyltransferase (Sobic.001G229000) that is present in 3 copies in the region (known in Arabidopsis as KATAMARI1/MUR3). KATAMARI1/MUR3 knockouts have shown disruption of the endomembrane organization and actin filaments, defects in cell elongation, a dwarfed phenotype, and a 50% reduction in fucose. It encodes a xyloglucan galactosyltransferase that interweaves with cellulose microfibrils in the cell wall (Madson *et al*., 2003; Tamura *et al*., 2005; Kong *et al*., 2015). The activity of the MUR3 protein was independently discovered to confer sensitivity to salt stress (Li *et al*., 2013). Additionally, a GDSL-like lipase (Sobic.001G228100) 48 kbp from the SNP had the second highest prediction score and this gene has been confirmed to cause mutations in epicuticular wax deposition in sorghum *bloomless* mutants (Jiao *et al*., 2018). A thick epi-cuticular wax layer forms a water vapor barrier in sorghum aerial organs which is thought to be key to adaptation for sorghum’s drought tolerance (Jordan *et al*., 1984).

The analysis of the second significant marker using the highest PlantCaduceus prediction score in this region, is a gene model of a lysine catabolism enzyme, hydroxyglutarate synthase with a 1338 domain-of-unknown-function (Sobic.001G227800; Supplementary Table 2). Lysine is the limiting essential amino acid in cereal crops and it is maintained in low concentrations in their tissue; Lysine synthesis is regulated by synthesis rate and catabolism (Stepansky *et al*., 2006). Overexpression of lysine-synthesizing enzymes leads to feedback inhibition and lowers agronomic yields (Yang *et al*., 2021). The intense feedback inhibition may be necessary to balance the carbon:nitrogen ratio to sustain high growth rates (Stepansky *et al*., 2006). Although not immediately obvious for a δ^13^C_leaf_ candidate gene, lysine-synthesizing mutants indeed demonstrate reduced photosynthetic rates, lower stomatal conductance, reduced starch content, and complex metabolic reprogramming that could suggest it is related to the δ^13^C_leaf_ phenotype and a reflection of *WUE*_i_ (Cavalcanti *et al*., 2018). Neighboring genes in linkage disequilibrium may explain how high-lysine feedback inhibition reduces photosynthesis and growth: Plastid Transcriptionally Active Chromosome 3 (pTAC3; Sobic.001G228300) is a plant nuclear encoded gene that activates and exerts nucleus control over chloroplast transcription activity of photosynthetic machinery genes (Yagi *et al*., 2012). There is a precedent for this although at a different genomic region. An extremely low δ^13^C_leaf_ mutant in sorghum was discovered and genetic analysis concluded that Chloroplast Import Apparatus 2 (CIA2) was most likely the causal gene (Sobic.010G250100) (Rizal *et al*., 2017). CIA2 encodes a transcription factor that regulates chloroplast translocon gene expression genes and thus the chloroplast import of nuclear-encoded proteins (Sun *et al*., 2001). Taken together, the genes under this SNP might be a metabolic switch that is responsive to water-stress or an imbalance between carbon and nitrogen. We hypothesize that a plant triggered by high/low lysine responds by i) simultaneously reinvesting captured carbon into epicuticular wax deposition to prevent further water loss from leaves, ii) initiating the nucleus to signal to the chloroplast to down-regulate photosynthetic apparatus. Future work characterizing the physiology of genetic knockouts lines for these genes in conjunction with δ^13^C_leaf_ will further the understanding of δ^13^C_leaf_ and *WUE*_i_ in C_4_ plants.

## Conclusions

The phenotype of δ^13^C_leaf_ has the potential to be a high-throughput proxy for *WUE*_i_ in C_4_ species; however, this can be confounded by both by genetic and environmental factors. Most estimates of δ^13^C_leaf_ heritability within C_4_ species have been based on small populations which can minimize the genetic and phenotypic variation. This work presented a survey of δ^13^C_leaf_ across a phenotypically and genetically diverse population of sorghum grown under field conditions. Heritability of δ^13^C_leaf_ was determined to be 57.5% and provides evidence that selection within breeding populations could potentially alter δ^13^C_leaf_. Genetic variation in δ^13^C_leaf_ can be statistically associated with specific genes and identifying their function will provide new insights on future targets to manipulate and explore δ^13^C_leaf_’s relationship to *WUE*_i_, the CCM, and post-photosynthetic fractionation in C_4_ plants.

The following supplementary data are available at JXB online:

Table S1. Gene models, annotations, and fine-mapping metrics for QTL on chromosome 9.

Table S2. Gene models, annotations, and fine-mapping metrics for QTL on chromosome 1.

## Acknowledgements

We wish to thank Dr. Stephen Kresovitch and Dr. Zachary Brenton for kindly providing access to the sorghum lines and experiments in this study and the personnel in his laboratory for helping with sampling plant material. This work was funded by a grant from Department of Energy (DE-SC0014395) to ABC and DPS.

